# Wampa is a dynein subunit required for axonemal assembly and male fertility in *Drosophila*

**DOI:** 10.1101/668012

**Authors:** Elisabeth Bauerly, Kexi Yi, Matthew C. Gibson

## Abstract

Axonemal dyneins are motor proteins that form the inner and outer arms of the axoneme in cilia and flagella. Defects in dynein arms are the leading cause of primary ciliary dyskinesia (PCD), which is characterized by chronic respiratory infections, situs inversus, and sterility. Despite current understanding of pathological features associated with PCD, many of their causative genes still remain elusive. Here we analyze genetic requirements for *wampa* (*wam*), a previously uncharacterized component of the outer dynein arm that is essential for male fertility. In addition to a role in outer dynein arm formation, we uncovered additional requirements during spermatogenesis, including regulation of remodeling events for the mitochondria and the nucleus. Due to the conserved nature of axonemal dyneins and their essential role in both PCD and fertility, this study advances our understanding of the pathology of PCD, as well as the functional role of dyneins in axonemal formation and spermatogenesis.

## Introduction

Cilia and flagella represent evolutionarily ancient organelles and their components are highly conserved across eukaryotes. In most motile cilia, hundreds of proteins collectively form an interior structure, known as the axoneme, which is assembled into a stereotyped 9+2 arrangement of microtubules [1]. Due to the overwhelming abundance of cilia within diverse animal cell types, disruption of axonemal assembly can lead to severe and pleiotropic defects. In humans, these defects are collectively referred to as ciliopathies or primary ciliary dyskinesia (PCD) for ciliopathies associated with motile cilia. Chronic respiratory issues, congenital heart defects, situs inversus, and sterility are some of most common phenotypes resulting from PCD and can be found singularly or in various combinations, depending on the genes affected [2, 3]. In PCD patients, up to 76% of affected males exhibit reduced sperm motility, resulting in decreased fertility [4]. Disruption in the axonemal outer dynein arms is the primary cause of PCD and is found in nearly 80% of all diagnosed patients [5, 6]. However, to date only about 30 different genes have been associated with PCD. There are cases, such as that of Joubert’s syndrome, where only 50% of the causative genes have been identified [7, 8]. Due to the complexity of the axonemal assembly, determining additional genes essential for the proper formation of cilia is a daunting yet essential task [7, 9].

Some of the challenges in elucidating causative genes associated with PCD come from the commonality of the phenotypes, which can lead to misdiagnoses [2, 3, 8]. Currently, visible axonemal abnormalities are the most reliable way of diagnosing PCD, but many disease-associated alleles could affect function without affecting structure. These difficulties are exemplified by dyneins, which affect transportation of proteins critical for ciliogenesis. Dynein disruption can lead to defective ciliogenesis yet do not always disrupt overall architecture. *Lis-1* is an example of one such dynein that performs multiple functions in multiple tissues. In humans, loss of Lis1 causes a smoothening of the cerebral tissue and disrupted neurons resulting in severe developmental abnormalities. In addition to its role in neurogenesis, Lis1 also plays essential roles during both oogenesis and spermatogenesis [10, 11]. Analysis of both *Drosophila* and *Chlamydomonas* models uncovered several novel roles for Lis1 during spermatogenesis, such as regulating centrosome positioning and facilitating attachment between the nebenkern, basal body, and nucleus without disrupting the architecture of the axoneme. These defects are believed to be caused by a disruption in dynein function, and thus Lis1 may have a yet-unrecognized role in PCD [12, 13]. However, since *Drosophila* is known to not utilize intra-flagellar transportation, the mechanism of transport during the assembly of the flagellum, and thus, consequences of its disruption, remains unknown.

In *Drosophila,* sensory neurons and sperm are the only ciliated cell types, which facilitates loss-of-function genetic analysis of axonemal assembly without the confounding effects of lethality or pleiotropic phenotypes. Here, using *Drosophila*, we characterize a previously unidentified component of the axoneme, *CG17083,* which we named *wampa (wam).* By generating mutant alleles of *wam* using CRISPR/CAS9 targeted mutagenesis, we demonstrate that it is essential for the formation of the outer dynein arms of the axoneme in the developing sperm flagella. Consequently, full loss of flagellar motility was also observed. In addition to the loss of the outer dynein arms, *wam*^*KO*^ homozygous males were fully sterile and displayed several unexpected defects during spermiogenesis, including mis-localization of mitochondria and malformation of the nuclear head. These phenotypes are strikingly similar to defects observed in mammalian models that disrupt intra-manchette transportation during spermatogenesis, suggesting a similar transport mechanism may operate during *Drosophila* spermatogenesis.

## Results

### *Wam* is essential for male fertility

Spermatogenesis in *Drosophila* begins at the apical tip of the testes, where the germline stem cells reside. Approximately every 10 hours, these stem cells asymmetrically divide to produce two daughter cells, one of which will remain in contact with the hub cells in the niche, while the other will move away in order to differentiate into an immature spermatogonia. This spermatogonia will undergo four rounds of mitotic divisions and, after an extended growth phase, will undergo two rounds of meiotic divisions to produce 64 interconnected spermatids (Figure 1A). Upon the completion of meiosis, the spermatid will then enter a phase known as spermiogenesis, which consists of 3 parts: The formation of the axoneme, drastic morphological re-shaping of the mitochondria, and re-modeling of the nucleus. These morphological steps are all necessary in order to finalize the production of the mature sperm. During this phase, hundreds of mitochondria aggregate around the basal body, which is embedded into the nuclear membrane. The mitochondria then fuse to form two separate masses that intertwine around one another forming one large sphere. The resulting structure is referred to as a nebenkern [14]. As the axoneme begins to elongate, the mitochondrial derivatives will unfurl from one another and grow with the developing axoneme, supporting it during an extreme elongation phase [15]. During axonemal elongation, the nuclear head also undergoes a dramatic reduction in size to nearly 200x smaller than it was at the beginning of spermiogenesis. At the end of the elongation and nuclear compaction stages, an actin complex individualizes all 64 sperm and cytoplastic debris is removed, resulting in fully mature, individualized, and highly coiled spermatozoa that can be transferred to the seminal vesicle (Figure 1B).

**Figure 1.**
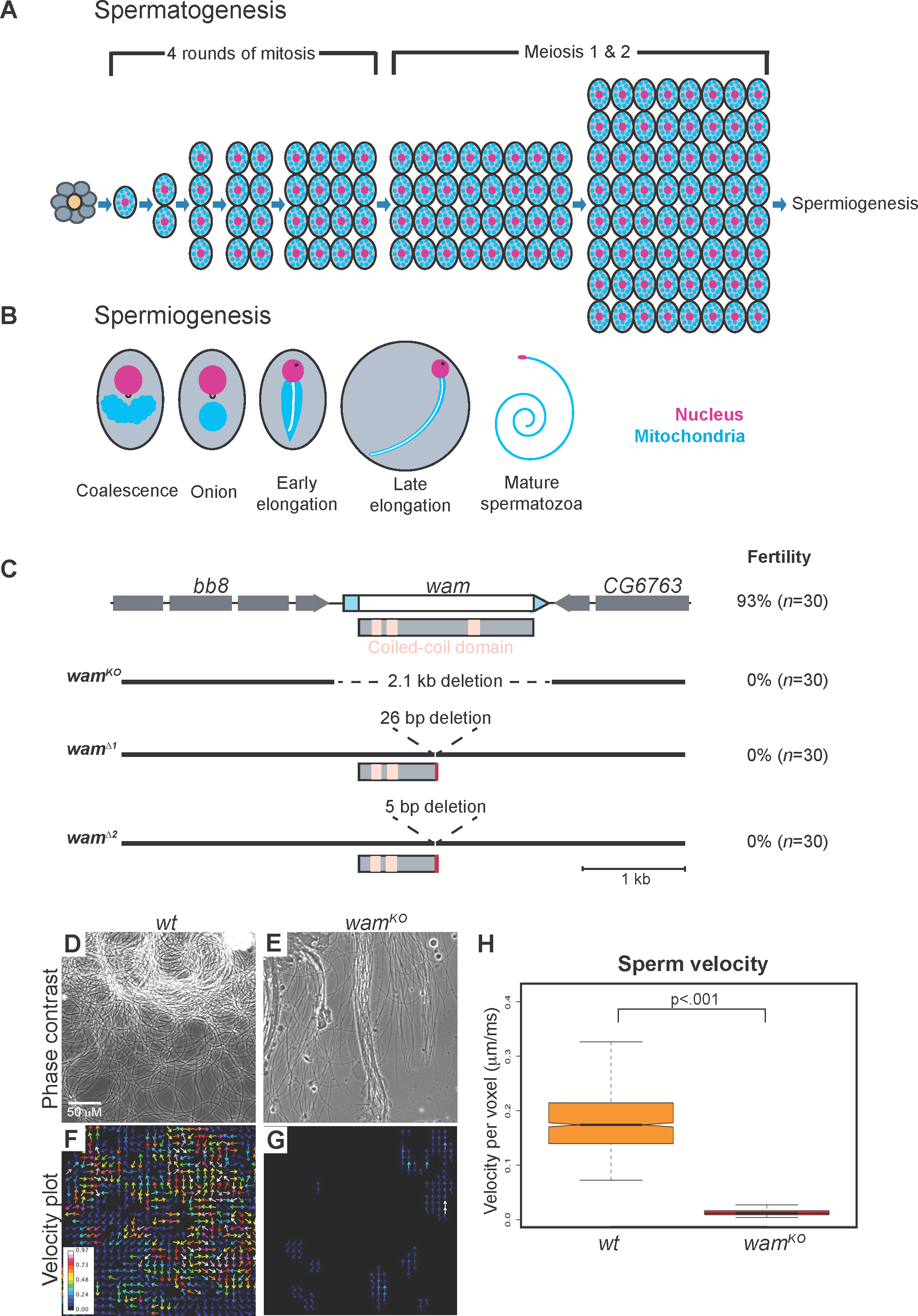
Wampa is essential for male fertility. (A) Schematic of spermatogenesis through the development of spermatids (adapted from Hales and Fuller, 1997) [38]. The germline stem cells surround the hub cells (*orange*) at the apical tip. The stem cells produce both a new stem cell as well as a gonialblast (*grey*). The gonialblast will undergo 4 rounds of mitotic divisions, followed by two rounds of meiosis, resulting in 64 spermatids. (B) Spermiogenesis. Immediately following meiosis, mitochondria coalesce around the basal body, which is embedded in the nuclear membrane, and fuse to form a large sphere-like structure known as the nebenkern. By the onion stage, the basal body is visible on the nucleus and axonemal assembly has initiated. As elongation proceeds, a phase-dark protein spot becomes apparent on the nucleus, the mitochondrial derivatives elongate with and support the developing axoneme (white line on mitochondria), and nuclear remodeling begins to compact the nucleus. The final steps in spermiogenesis will individualize and coil the sperm, forming mature spermatozoa. (C) Graphic of the *wampa* locus and CRISPR-generated alleles. CDR and UTRs are represented by white and blue boxes, respectively. The three coiled-coil domains are denoted by pink boxes on the protein schematic. *wam*^*KO*^ contains a large 2.1kb deletion that removes the entire gene region, while *wam*^*Δ1*^and *wam*^*Δ2*^ both generated small indels, which resulted in premature stop codons. Fertility quantification is labeled on the right of the schematic for each genotype. See also Figure S1. (D-G) Representative frame from a movie of live *wildtype* (D) and *wam*^*KO/KO*^ (E) sperm and a corresponding velocity heat map displaying actively moving sperm in *wildtype* denoted by warmer colors (F) and immotile sperm in *wam*^*KO/KO*^ denoted by cooler or lack of color (G). See also movie S1. (H) Box plot of the average sperm velocity per pixel, for both *wildtype* (orange, *n=*12 animals) and *wam*^*KO/KO*^ (red, *n=*10 animals) showing a lack of movement in sperm from *wam*^*KO/KO*^ males compared to *wildtype.*

*Wam* is a highly conserved gene that was predicted to be involved in outer dynein arm formation based on sequence similarity with *Chlamydomonas* [16, 17]. In order to assess its role in *Drosophila*, we employed CRISPR/CAS9 mutagenesis to create three mutant alleles. *Wam* encodes a 550 amino acid (aa) protein that contains three coiled-coiled domains (Figure 1C). We first generated a novel null allele, *wam*^*KO*^, by utilizing two gRNAs targeting either sides of the coding region, removing the entire gene locus. Additional alleles, *wam*^*Δ1*^ and *wam*^*Δ2*^, were generated using a single gRNA to target the middle of the coding region, which resulted in a truncated protein at 250aa that lacks the last coiled-coiled domain (Figure 1C and Figure S1). Phenotypically, homozygous animals from each allele appeared normal and displayed no obvious defects. However, when assayed for fertility, homozygous mutant males exhibited complete sterility while females produced similar amounts of progeny as *wildtype* (Figure 1C). All three *wam* alleles resulted in identical phenotypes in every assay and did not show any apparent defects in sensory behavior. For simplicity, only *wam*^*KO*^ is utilized from here on. Using both complementation and rescue experiments, we confirmed that the sterility was due to the loss of *wam* and not *bb8*, a neighboring gene which is also required for male fertility (Figure S2) [18]. Taken together, these data indicate that *wam* is required for male fertility.

### *Wam* is required for axonemal assembly

To investigate the cause of sterility in *wam*^*KO*^ homozygous males, we first examined the testes using phase contrast microscopy and discovered that sperm exhibited a complete loss of flagellar motility (Figure 2D-2H and movie S1). We also observed that the seminal vesicles of *wam*^*KO/KO*^ animals were significantly reduced in size compared to *wildtype*, indicative of a defect in spermatogenesis prior to sperm loading. To quantify this observation and examine the phenotypic progression, we held males as virgins for two weeks and examined the morphology of the testes. At day one post-eclosion, the gross morphology of testes and associated structures were indistinguishable between *wam*^*KO/KO*^ and *wildtype* animals (Figure 2A,D). However, as the animals aged, the seminal vesicle of *wam*^*KO/KO*^ animals failed to fill with sperm and increase in size as observed in *wildtype* (Figure 2, *arrows*). Instead *wam* mutant animals showed an enlargement of the basal end of the testes (Figure 2C and E-F, arrow heads), further consistent with a failure of sperm to enter the seminal vesicles.

**Figure 2.**
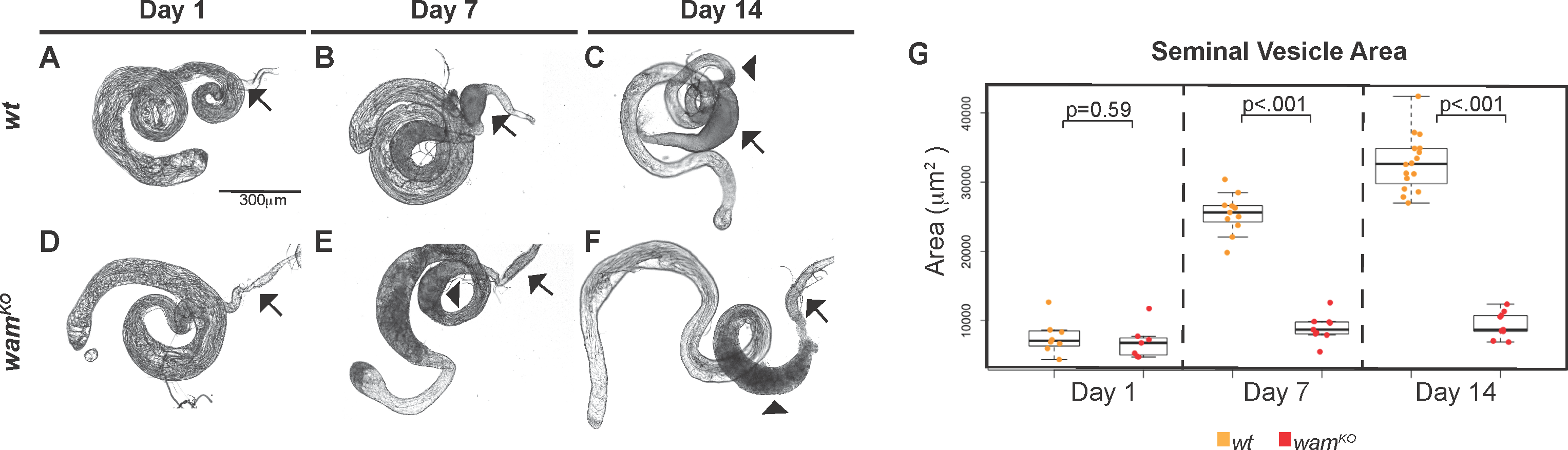
Sperm from *wampa* mutants are unable to enter the seminal vesicle. (A-F) *Wildtype* (A-C) and *wam*^*KO*^ (D-F) homozygous males were held as virgins for two weeks and their testes were examined for morphological differences during this time. At day 0, testes from *wildtype* (A) and *wam*^*KO*^ (D) are comparable. However, by day 7, the differences in the seminal vesicles can be detected as *wildtype* seminal vesicles begin to enlarge (B, *arrow*), whereas *wam*^*KO*^ remains approximately the same size (E, *arrow*). At day 14, seminal vesicle size remains the only difference observed (C and F, *arrows*), indicating a failure for sperm to be loaded and, as a result, the basal end of the testes becomes enlarged in *wam*^*KO*^ (E and F, *arrowhead*) compared to *wildtype* (C, *arrowhead*). (G) Box plot of the average area of the seminal vesicles for both *wildtype* and *wam*^*KO*^ homozygotes at day 0 (*wt, n=*8 and *wam*^*KO*^, *n*=7), day 7 (*wt, n*=12 and *wam*^*KO*^, *n*=10), and day 14 (*wt, n*=17 and *wam*^*KO*^, *n*=10). *wildtype* seminal vesicles gradually increase in size over time, while *wam*^*KO*^ seminal vesicles stay approximately the same size.

Since *wam*^*KO/KO*^ mutants displayed complete immotility of the sperm flagellum, we next examined the ultrastructure of the axoneme. In normal *wildtype* sperm, outer dynein arms are an essential axonemal substructure known to be required for motility [9] (Figure 3A,B). Strikingly, transmission electron microscopy (TEM) revealed that *wam*^*KO/KO*^ sperm flagella lacked outer dynein arms along the length of the axoneme (Figure 3A-C). We next sought to determine whether *wam* functions as a cytoplasmic dynein involved in the pre-assembly transport of the outer dynein arms, such as *Wdr92*, or whether it is directly incorporated onto the mature flagellum [19]. To resolve this issue, we generated a transgenic line expressing Wam::mScarlet-I under the control of the endogenous *wam* promoter. Analysis of this tagged construct revealed that *wam* is in fact incorporated into the axoneme and can be visualized along the entire length (Figure 3D). To test functionality of the Wam::mScarlet-I construct, we performed a rescue assay in wam homozygous males. The fusion protein was able to fully restore fertility to *wildtype* levels (Figure S2), consistent with the interpretation that *wampa* is an essential component of the sperm flagellar axoneme that is required for the assembly of the outer dynein arms.

**Figure 3.**
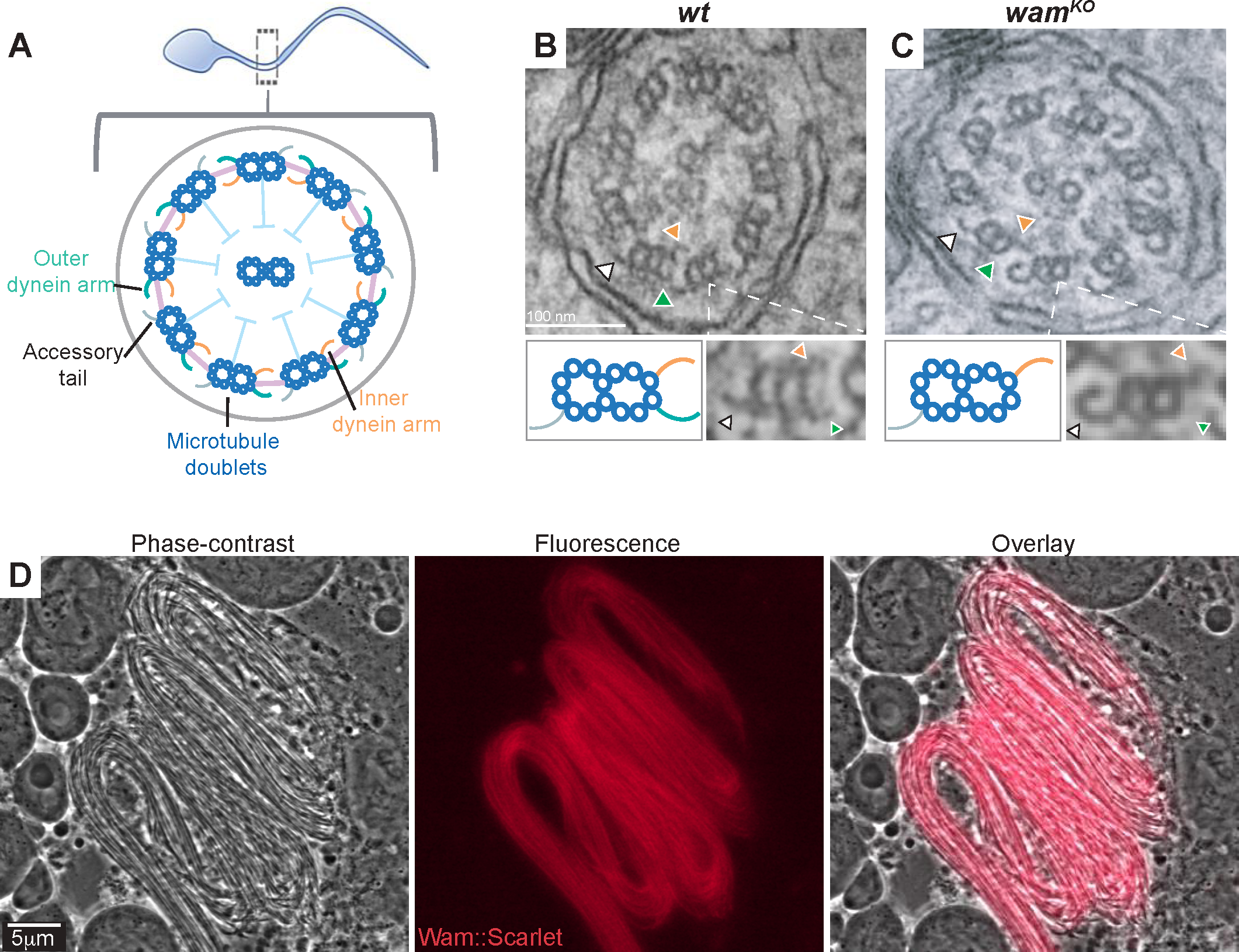
Wampa is essential for the assembly of the outer dynein arms. (A-C) Schematic of the flagellar axoneme (A). TEM images of a *wildtype* axoneme showing the inner (*orange arrow*) and outer (*green arrow*) dynein arms on the microtubule doublet, opposite from the position of the accessory tail (*white arrow*) (B). Outer dynein arms are notably absent in *wam*^*KO*^ homozygotes (C). A blowup of an individual doublet is displayed below the EM images on the right, with a corresponding cartoon on the left. (D) Live phase contrast and fluorescent images from testes expressing the Wam::mScarlet-I tag, demonstrating endogenous Wam localization along the entire length of the flagella of a developing spermatid bundle.

### Mitochondrial localization and elongation defects are observed in *wam*^*KO*^

In addition to the lack of flagellar motility, we observed that sperm failed to fully individualize and coil in *wam*^*KO/KO*^ testes (Figure 1D-E). Failure to complete these last steps of spermiogenesis could be an indicator of defects in earlier stages, prior to spermiogenesis [20]. To confirm that *wam* was expressed prior to axonemal assembly, we analyzed expression of the Wam::mScarlet-I reporter construct in adult testes. In premeiotic cells, *wam* accumulated in the cytoplasm and was excluded from the nucleus (Figure 4a). During meiosis, *wam* was diffuse throughout the cell but exhibited increased aggregation in puncta decorating the spindle poles (Figure 4B,C). By the onion stage, *wam* localization began to concentrate around the nucleus and during early elongation accumulated in a transient microtubule-based structure known as the manchette (Figure 4D-E’). The manchette resides on top of the nuclear head during the remodeling process (Figure 4F) and is known to be involved in transportation of proteins for critical for spermiogenesis in mammals.

**Figure 4.**
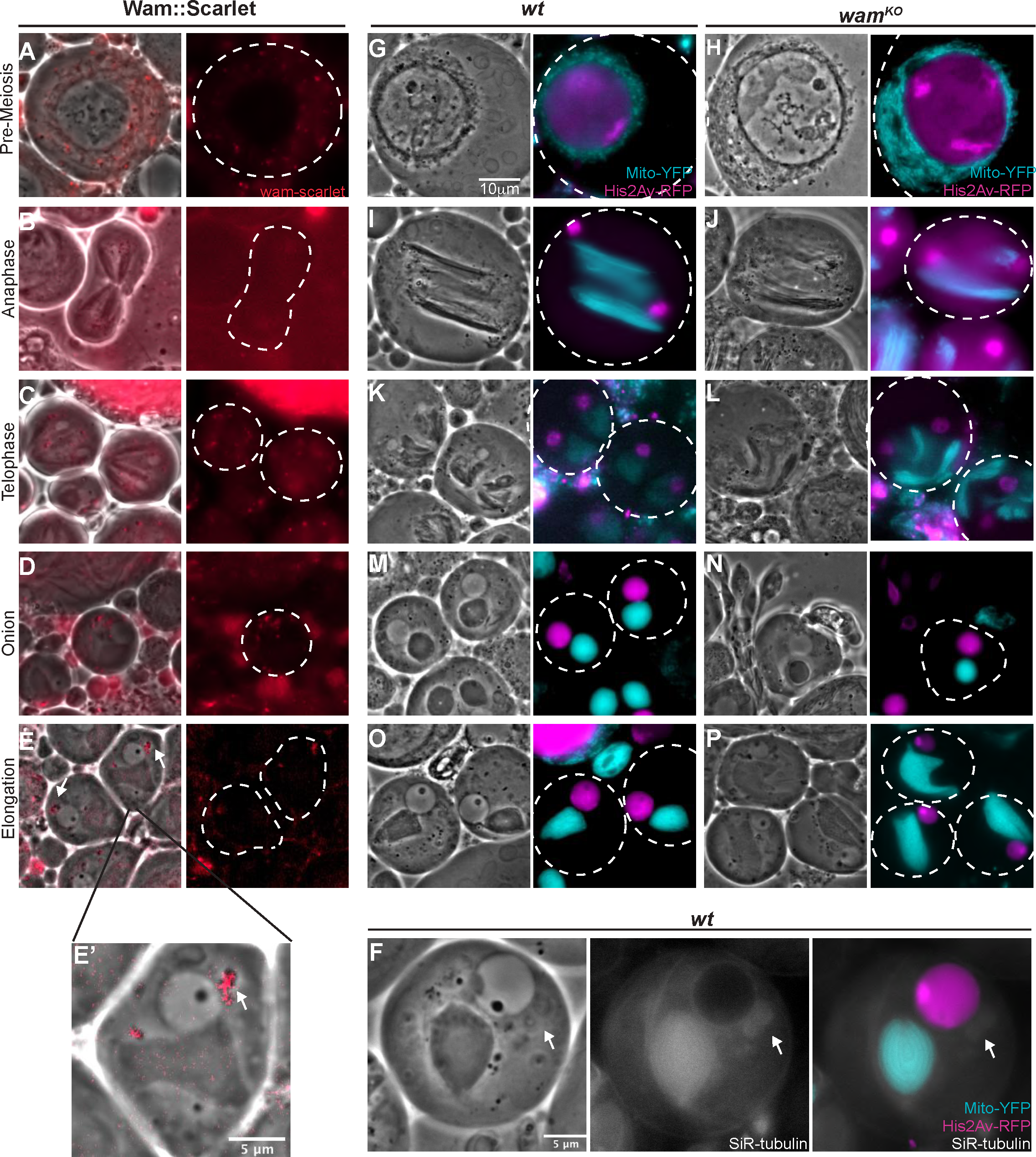
Wam is required for mitochondrial localization during meiosis. (A-E’) Merged phase-contrast and epifluorescent images showing the localization of Wam::mScarlet-I during chronological stages of spermatogenesis. Prior to spermiogenesis, *wam* is diffuse throughout the cytoplasm (A-C). During meiosis, there is a small accretion near the spindle poles and around the mitochondria (B-C). At the onion stage, Wam accumulates around the nucleus (D) and during early elongation, Wam is localized in the phase dark manchette on the nuclear head (E,E’). (F) Phase-contrast (*left*) and fluorescent (*middle and right*) images showing localization of the manchette (*white arrow*) relative to the nucleus in a *wildtype* spermatid. (G-P) Phase-contrast (*left*) and fluorescent (*right*) images of chronological stages of spermatogenesis. In the pre-meiotic stages, mitochondria can been seen surrounding the nucleus for both *wildtype* (G) and *wam*^*KO/KO*^ (H). In spermatocytes from *wildtype* males, mitochondria are evenly distributed (I and K). However, in *wam*^*KO*^ sperm, the distribution of mitochondria is perturbed during meiosis (J and L). By the onion stage, most of the mitochondria are packed into the expected nebenkern and appear superficially normal through elongation (M-P).

To examine whether *wam* affected these earlier of spermatogenesis, we fluorescently labeled both the DNA and mitochondria and, in conjunction with phase contrast imaging, analyzed various stages of spermatogenesis. The early, pre-meiotic stages of spermatogenesis appeared unperturbed (Figure 4G,H). However, starting in meiosis, we observed mitochondrial aggregation defects. During meiosis, *wildtype* mitochondria align evenly along the spindle in order to ensure equal segregation during cytokinesis (Figure 4I,K). In contrast, mitochondria failed to appropriately line up along the spindle in a subset of cells in *wam*^*KO/KO*^ mutants (Figure 4J,L). By the onion stage and through elongation (Figure 1B) most of the cells in *wam*^*KO/KO*^ testes appeared superficially normal (Figure 4M-P), but more detailed analysis revealed a range of additional defects.

Following the meiotic divisions, mitochondria associated with each developing spermatid aggregate around the basal body and are then fused into the two separate derivatives that form the nebenkern [14]. While the majority of onion- and elongation-stage cells appeared normal in *wam*^*KO/KO*^ mutants, a small subset displayed a fragmentation phenotype where small clusters of unincorporated mitochondria were observed adjacent to the nebenkern (Figure 5A-D, *arrows*). TEM analysis revealed a surprising defect with the mitochondrial derivatives not obvious by brightfield microscopy. At the beginning of the elongation phase, the two mitochondrial derivatives that form the nebenkern normally unfurl from one another and extend along the growing axoneme, providing both physical and metabolic support [21]. In *wam*^*KO/KO*^ testes, the mitochondrial derivatives were consistently and significantly smaller than in *wildtype* (Figure 5E-G, arrows). As the mitochondria provide structural support for the developing axoneme and are essential for the formation of the flagellum, a disruption in their elongation could further destabilize the developing axoneme.

**Figure 5.**
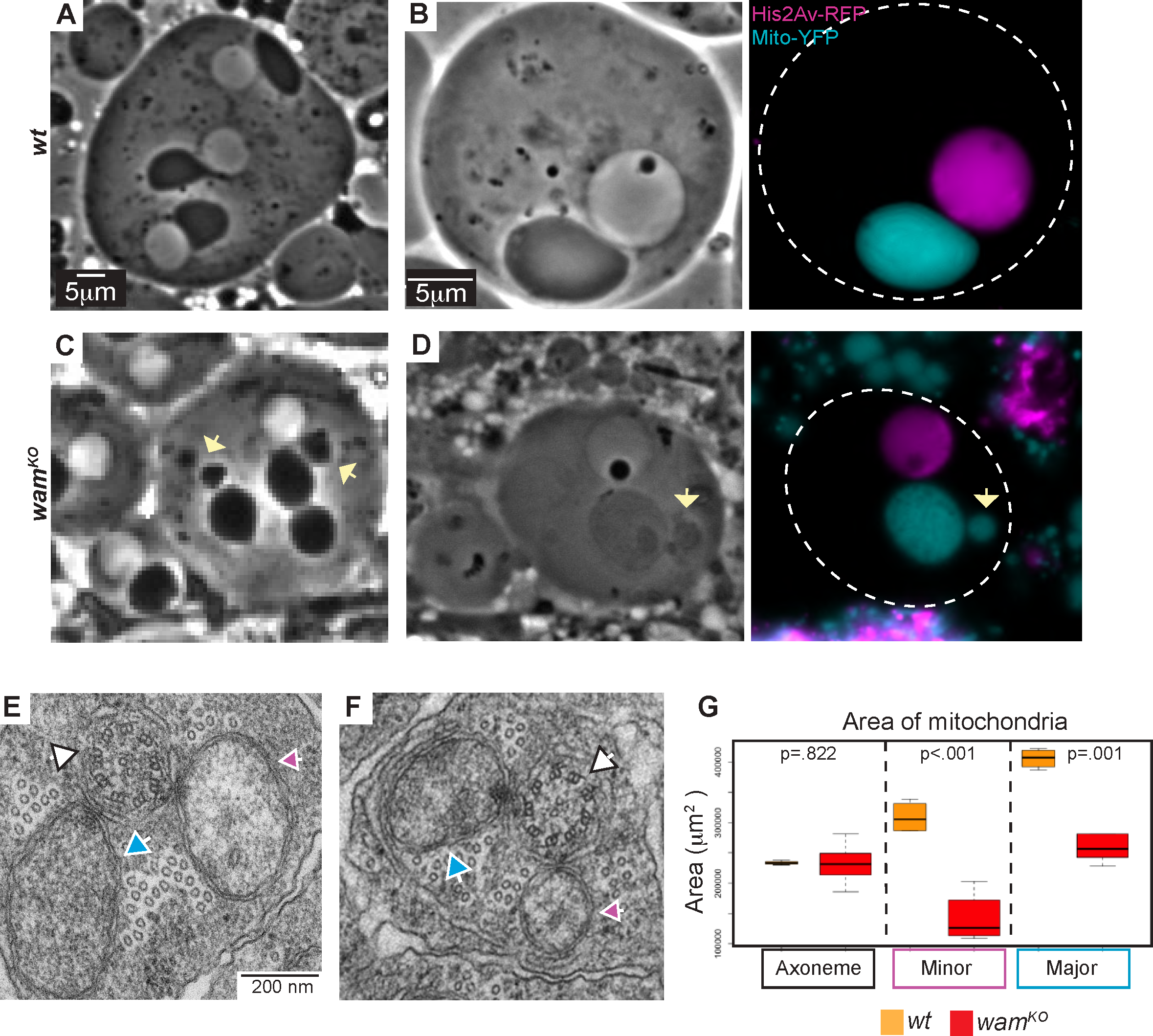
Mitochondrial elongation is stunted in *wam*^*KO*^. (A-D) Phase contrast and fluorescent images of onion stage spermatids from *wildtype* and *wam*^*KO/KO*^ testes at 20x (A and C) and 100x (B and D). Fragmentation of the nebenkerne can occasionally be observed in *wam*^*KO*^ spermatids (C and D, *yellow arrow*). (E-F) EM images of spermatids during elongation for both *wildtype* (E) and *wam*^*KO/KO*^ (F). The axoneme (*white arrow*) is visible with major (*blue arrow*) and minor (*pink arrow*) mitochondrial derivatives on either side. (G) Quantification of the area of the major and minor mitochondrial derivatives and axonemes for *wildtype* (*orange, n=*5) and *wam*^*KO/KO*^ spermatids (*red, n=*11). The axoneme is approximately the same size for both *wildtype* and *wam*^*KO/KO*^, but the major and the minor mitochondrial derivatives in *wam*^*KO/KO*^ show a severe decrease in size compared to *wildtype*.

### *wam* is required for nuclear remodeling

During spermiogenesis, every cell within a 64-cell cyst should contain a nucleus and a nebenkern. This ratio is always 1:1 and deviations from the expected ratio are typically due to meiotic defects, most commonly during cytokinesis [21, 22]. In *wam*^*KO/KO*^ testes examined by phase contrast microscopy, numerous onion-stage cells appeared to lack a nucleus, containing only the nebenkern. To examine these onion stage nuclei more precisely we used a transgenic line with a fluorescent DNA label. Unexpectedly, rather than a skewed ratio of nuclei to nebenkern, *wam*^*KO/KO*^ cells presented with severe defects in nuclear morphology, suggesting errors during nuclear remodeling (Figure 6A-D). Indeed, analysis of over 100 onion staged nuclei from both *wildtype* and *wam*^*KO*/KO^ testes revealed that even nuclei that appeared superficially normal were in fact slightly smaller than their *wildtype* counterparts (Figure S3). This observation is reminiscent of intra-manchette transportation defects that have been observed in mammals wherein defects in flagellar assembly correlate with defects in the reshaping of the nuclear head. Our imaging confirms the presence of the manchette on the nucleus of developing spermatids in both *wildtype* and *wam*^*KO*^ homozygous males (Figure 6 E,F). Together with the localization data (Figure 4E), these results indicate that *wampa* is required for intra-manchette transportation during spermatogenesis [23, 24].

**Figure 6.**
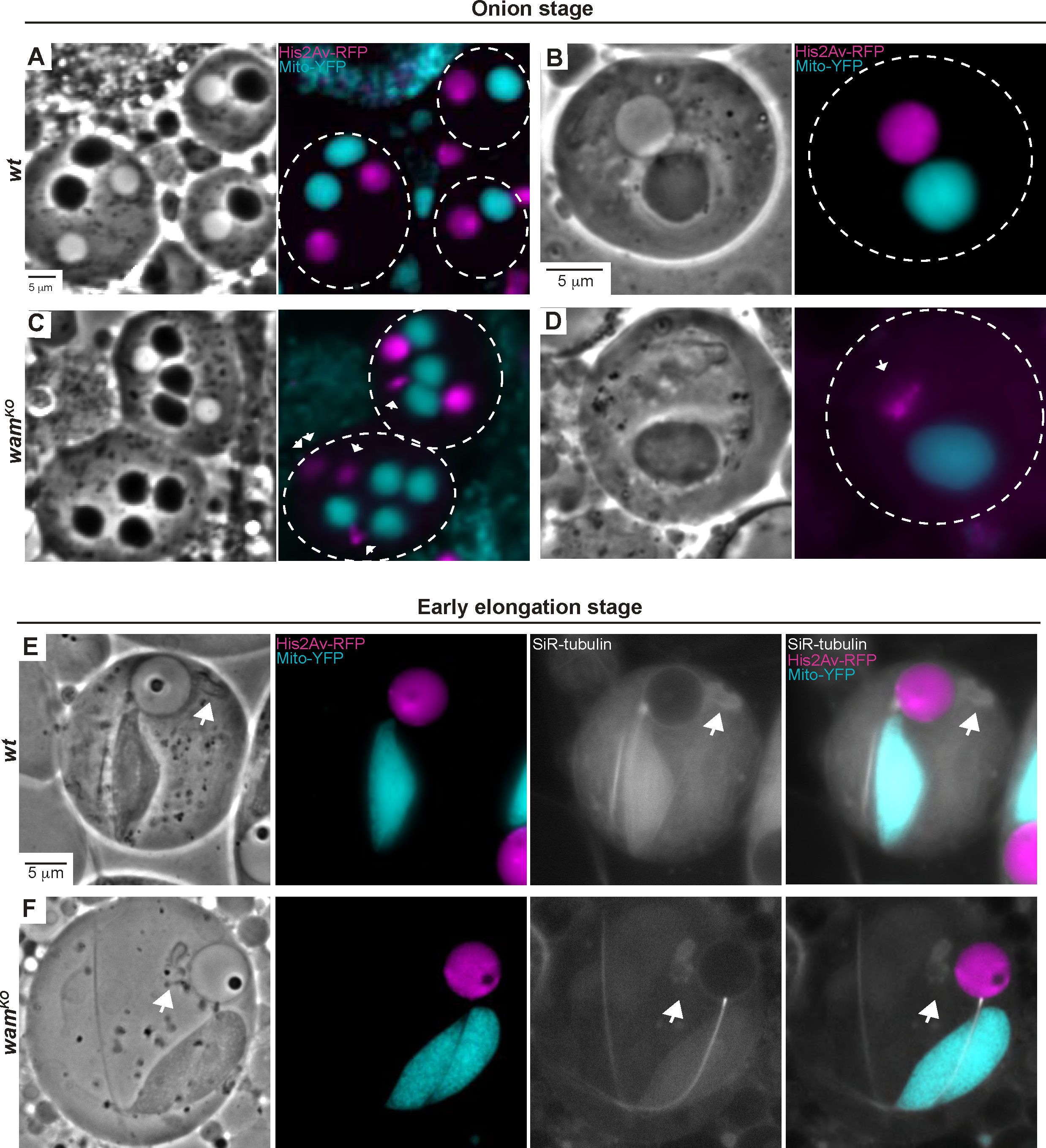
Wam regulates shaping of the nuclear head. (A-D) Phase-contrast and florescent images of onion stage spermatids from *wildtype* (A and B) and *wam*^*KO/KO*^ testes (C and D) shown at 20x and 100x, respectively. *Wildtype* cells display the expected 1:1 ratio of nebenkern to nucleus (A and B), whereas a corresponding nucleus can’t be seen by phase-contrast in *wam*^*KO/KO*^ for some cells. However, fluorescently-labeled DNA reveals the presence of nuclei which fail to properly condense through nuclear remodeling that occurs during spermiogenesis (C and D). (E-F) Phase-contrast and florescent images of spermatids during early elongation from *wildtype* (E) and *wam*^*KO/KO*^ males (F). SiR-tubulin allows microtubules to be visualized which reveal the presence of the manchette (*white arrow*). No differences are observed in these structures.

## Discussion

Ciliogenesis requires the coordinated activity of hundreds of different proteins to ensure proper form and function. Disruption of any one of these factors risks the potential of catastrophic failure of the tissues associated with the malformed cilia. Here we show that the *Drosophila* dynein-associated protein *wam* is essential for proper axonemal assembly as well as successful completion of spermatogenesis. In the absence of *wam* function, outer dynein arms were lost along the entire length of the axoneme, which was in turn correlated with defects in mitochondrial transport as well as shaping of the nuclear head. Since microtubules are broadly required during spermatogenesis and dyneins are well known to play critical roles in cell cycle progression [13], it is not surprising that dynein-related genes might have a profound effect on multiple different stages of spermatogenesis. However, the defects observed in *wam* are unique in that they show clear phenotypes in all three of the major processes occurring during spermiogenesis.

Our results indicate that *wam* is essential for proper formation of the flagellum and that when disrupted, outer dynein arms fail to assemble on the developing axoneme. The loss of the outer dynein arms along the axoneme leads to complete sperm immotility and sterility, demonstrating *wam’s* critical role in male fertility. Interestingly, *wam* has two mammalian orthologs, *CCDC114* and *CCDC63. CCDC114* has been identified in patients presenting with PCD related defects including chronic respiratory issues, heart defects, and situs inversus [16]. While fertility issues have not yet been reported in these patients, the authors did show that *CCDC114*, like *wam*, localizes along the entire length of the axoneme and is required for outer dynein arm assembly. Intriguingly, the other homolog, *CCDC63,* did result in male sterility when mutated in mice but, unexpectedly, retained the integrity of the outer dynein arms. The authors speculated that the retention of the outer dynein arms might be due to the compensation of *CCDC114* and suggested a double knockout would provide more insight on the function of *CCDC63* [25]. Nevertheless, while the outer dynein arms were intact in *CCDC63* mutants, spermiogenesis was associated with abnormal morphology of both the nuclear head and flagellar tail [25]. This is consistent with other defects we observed in *wam*^*KO/KO*^ testes and nicely demonstrates the conserved nature of the single copy of *Drosophila wam* and its ability to phenocopy defects observed in both mammalian orthologs.

In addition to its role in the formation of the axoneme, *wam* is also involved in mitochondrial remodeling. During our analysis of *wam* mutants, we observed that the mitochondria were not properly aligned along the meiotic spindle resulting in perturbation of their segregation during telophase through coalescence. This led to a small subset of nebenkerne that appeared to be fragmented during the onion stage. Interestingly, this phenotype bares a strong resemblance to phenotypes observed in *lis-1* mutants, which are thought to be a consequence of the disruption of dynein function [13]. Further TEM analysis also uncovered a defect in the mitochondrial derivatives associated with the elongating axoneme in *wam* mutants, which were consistently and significantly smaller than those observed in *wildtype*. This was an intriguing finding as it correlated with the observation of the fragmented nebenkerne, suggesting the possibility that not all of the mitochondria were properly incorporated into the mitochondrial derivatives, resulting in smaller derivatives at elongation. Still, it remains unclear why a larger percentage of nebenkerne did not appear fragmented. It is possible that the unfused fragments are in close enough proximity to the nebenkern so as to be indistinguishable or perhaps they are being degraded, resulting in less mitochondria forming the nebenkern and thus, smaller derivatives. It is also plausible that there is an unidentified defect which stunts the growth of the mitochondrial derivatives during elongation. While the precise cause of the elongation defect is not clear, the observation of fragmented nebenkerne and misaligned mitochondria during meiosis strongly suggests that *wam* is involved in mitochondrial localization during coalescence. It has previously been suggested that the movement of mitochondria during spermatogenesis is dependent on microtubules, and specifically on dynein, and disruption of dynein-associated genes have been shown to perturb the mitochondrial dynamics [13, 14, 26]. Indeed, our imaging reveals that the mitochondria and microtubules co-localize during spermatogenesis (Figure 4F and 6E,F). The disruption of the nebenkerne and of the elongation of the mitochondrial derivatives are strong indicators that *wam* is involved in the dynamic movement and localization of mitochondria during spermatogenesis.

Perhaps the most unexpected defect associated with *wam* alleles is the disruption in the nuclear shaping. Our data shows that the morphogenesis of the nuclear head was perturbed in *wam* mutants, resulting in condensation defects. In mammals, this nuclear re-shaping process is known to be facilitated by the manchette, a microtubule-based structure which links the developing spermatid to the cell surface through microtubule interactions [27, 28]. Defects in either the formation of the nuclear head or the flagellum can result in reciprocal abnormalities in the other process. This has been shown to be a result of disruptions in intra-manchette transportation of proteins essential for spermatogenesis in mammals [23, 29]. In fact, over a dozen genes in mammals have recently been identified to be involved in intra-manchette transportation that perturb both the remodeling of the nuclear head as well as the formation of the flagellum [30]. While there is still a lot unknown about the function of the manchette, it is suggested to be very similar to intraflagellar transportation [23]. Interestingly, we identify here a very similar structure in *Drosophila* (Figure 4F and 5E,F, white arrow) that has previously been referred to as the dense complex [26, 31, 32] and has been reported to be involved in the reshaping of the nuclear head [26, 33, 34]. However, this is the first time that is has been linked to disruption in both the formation of the nuclear head and the flagellum, as is reported in mammals. Given that intraflagellar transport has been shown to not be utilized during *Drosophila* spermatogenesis [35, 36], it is very likely that *Drosophila*, like mammals, requires intra-manchette transportation in order to move essential proteins to the nucleus, as well as to the axoneme and mitochondria during these remodeling events. This would explain the defects observed in *wam* mutants and strongly suggests that insects and mammals have a more similar approach to completing spermatogenesis than previously recognized.

In summary, here we identified Wam as an essential component of axonemal assembly and elucidated roles throughout spermatogenesis, which are reminiscent of a role during intra-manchette transportation. Disruption of both of the mammalian orthologs of *wam, CCDC63* and *CCDC114*, results in PCD associated with sterility, severe respiratory distress, heart malformations, and laterality defects. These syndromes are due to errors either in spermiogenesis that disrupt nuclear remodeling and the formation of the flagellum or in loss of the outer dynein arms during ciliogenesis. Since there are hundreds of genes affiliated with ciliogenesis, many PCD-related diseases are considered rare and often difficult to diagnose. Despite extensive effort, at least a third of genes that cause PCD remain elusive and those that are known do not always have a clear functional role [37]. Here we show that *wam* is able to recapitulate the phenotypes that result in PCD defects observed in both of its mammalian orthologs. As such, further analysis of *Drosophila wam* could be a productive avenue for investigating molecular mechanisms underlying the pathology of diseases associated with ciliogenesis.

## Acknowledgements

We thank the Bloomington Stock Center for fly stocks, Eric Hill for a critical reading of the manuscript, Steve Hoffman for assisting with microscopy, Brian Slaughter for aiding with the STICS analysis, Cindi Staber for teaching testes dissection and a critical reading of the manuscript, and Takuya Akiyama for Ubi-mCherry DNA, as well as extensive discussion and suggestions. This work was supported by funding from the Stowers Institute for Medical Research and NIH Grant R01-GM111733 to M.G..

## Author contributions

M.G. and E.B. designed the project. K.Y. acquired EM images. E.B. performed all experiments and analyzed all data. M.G. and E.B. wrote the manuscript.

## Supplemental information

**Figure S1.**
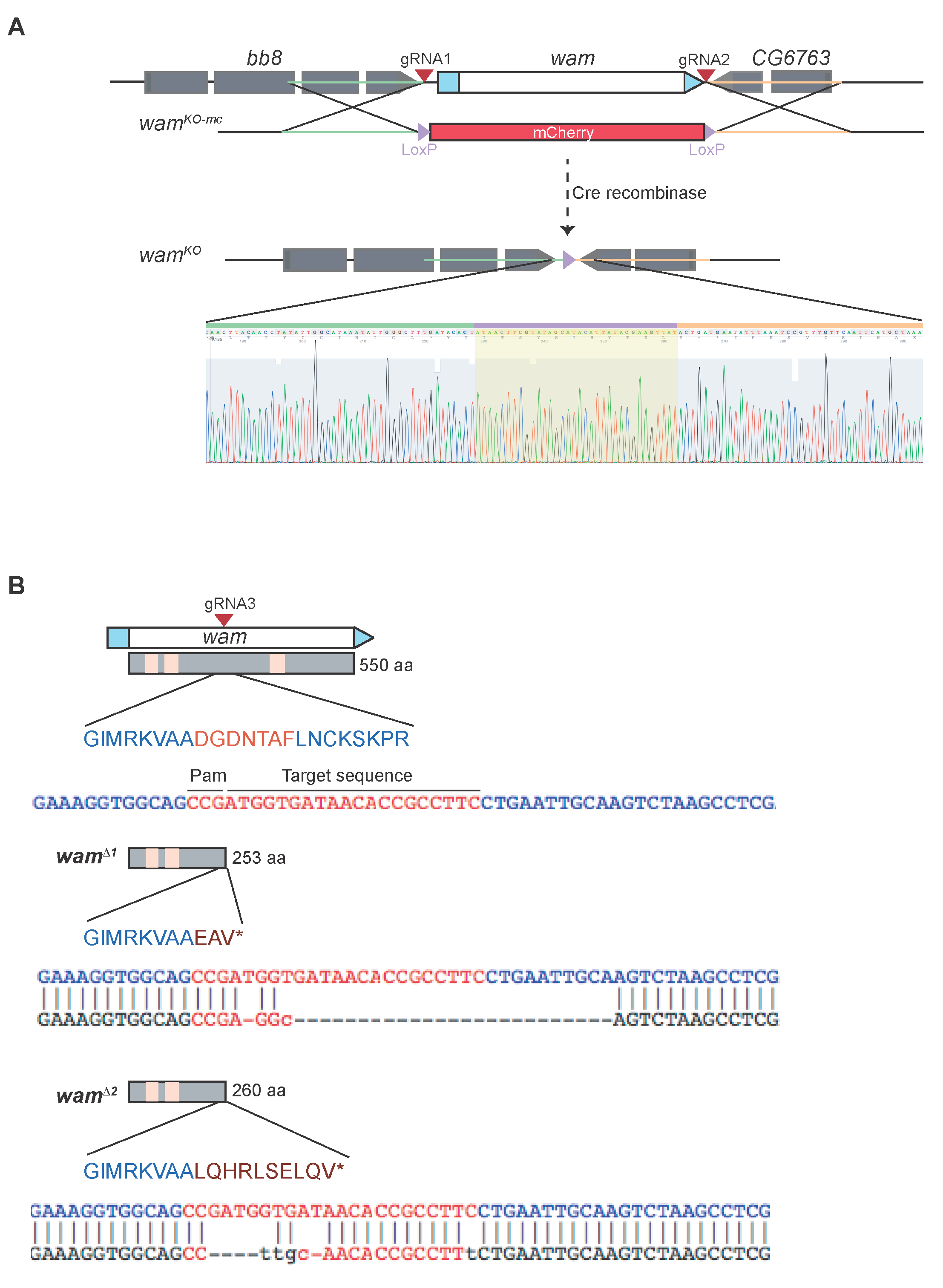
Generation of *wampa* alleles. (A) Schematic displaying general design and confirmation for the generation of *wam*^*KO*^ allele. Two gRNAs were used to create double strand breaks on either side of the *wampa* locus (*red arrows*). The breaks were repaired via homologous recombination from a donor template that contained 1 kb homology upstream (*green line*) and 1 kb homology downstream of *wampa* (*peach line*), as well as an mCherry cassette flanked by *loxP* sites (*purple arrows*). This generated the *wam*^*KO-mc*^ line. We then used CRE-LOX recombination to remove the mCherry cassette, generating the *wam*^*KO*^ line. This line was validated with sanger sequencing using primers in the 5’ and 3’ homology arms that resided outside of the gene region and showed a full removal of the entire coding region for *wampa*, leaving only the expected *loxP* site. (B) Schematic displaying the design and confirmation for the generation of *wam*^*Δ1*^and *wam*^*Δ2*^ alleles. A single gRNA was used to create breaks in the middle of the coding region for *wampa*. The break was repaired via non-homologous end joining. Due to the inefficiency of this repair process, indels were created generating two separate lines. Sequencing confirmed that *wam*^*Δ1*^ had indels that resulted in three extra amino acids and an approximately 26bp deletion, while *wam*^*Δ2*^ resulted in a smaller deletion of only 5bp with indels that generated 10 extra amino acids. Both alleles had an early stop site that terminated at 250aa, just under half way through the 550aa protein.

**Figure S2.**
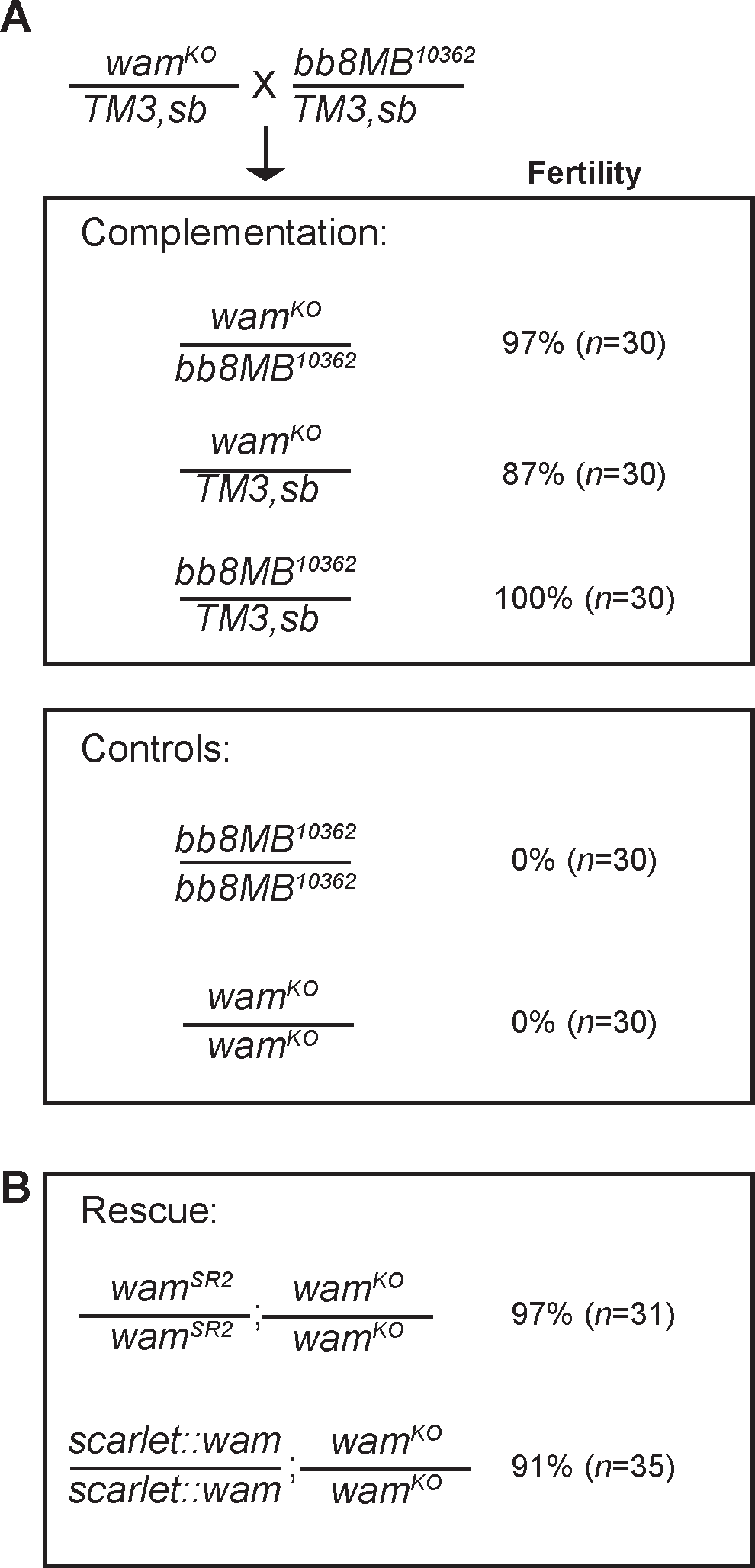
*wampa* is essential for male fertility. (A) Complementation tests were performed by generating trans-heterozygotes of *wam* and *bb8*. All progeny were collected and screened for fertility; homozygotes from the parental stocks were used for controls. Homozygous males from both *bb8*^*MB10362*^ and *wam*^*KO*^ are sterile as homozygotes, however, as trans-heterozygotes, they are fertile indicating that *bb8* is not contributing to the sterility observed in *wam*^*KO*^. (B) The rescue construct, *wam*^*SR2*^, and the tag construct, *Wam::mScarlet-I*, were both examined in the background of *wam*^*KO/KO*^ and tested for fertility. Both lines were functional and able to fully rescue the sterility defects observed in *wam*^*KO/*KO^ males.

**Figure S3.**
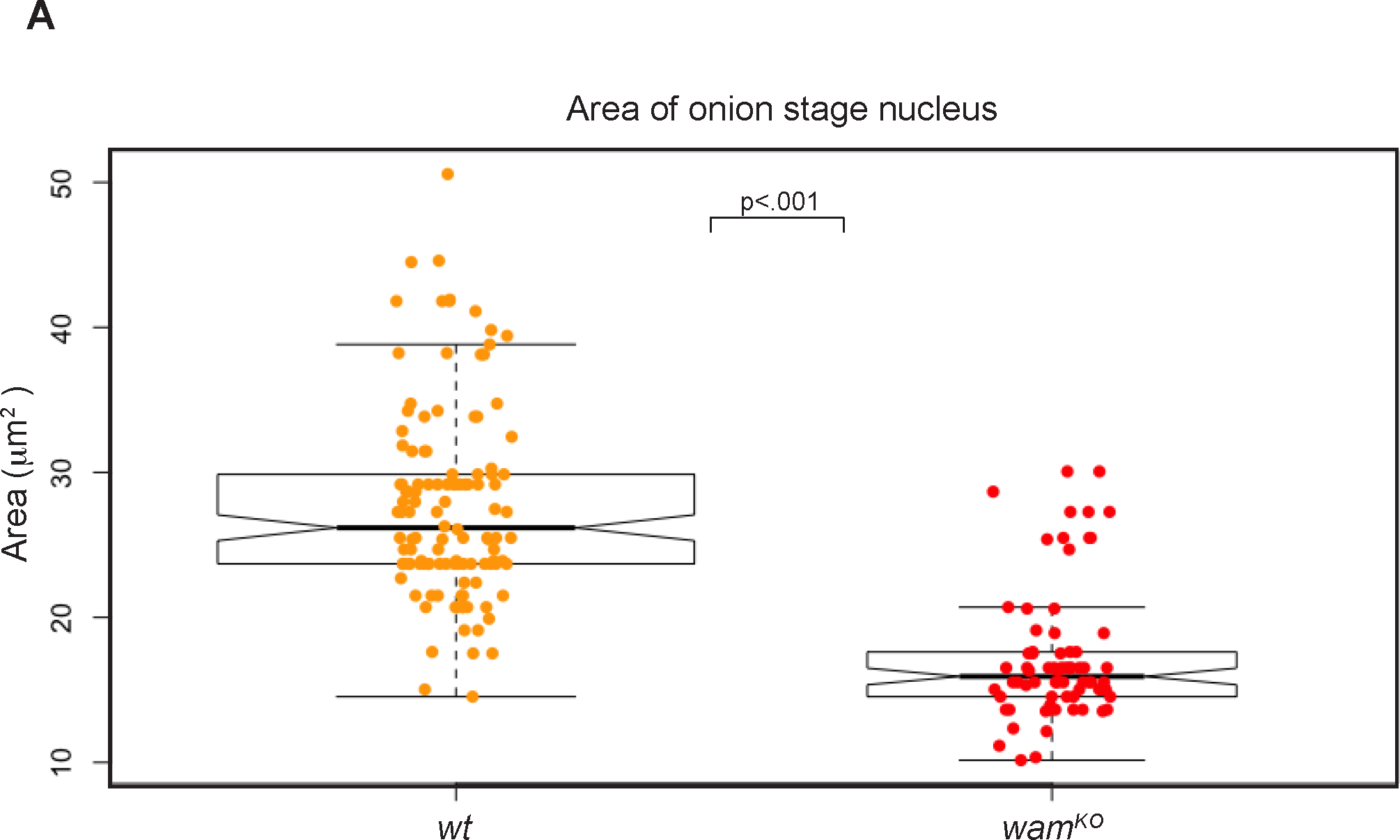
Wam regulates shaping of the nuclear head. (A) Quantification of nuclear area in developing spermatids at the onion stage for *wildtype* (orange, *n=*144) and *wam*^*KO/KO*^ (red, *n=*116). Nuclei were generally reduced in size in *wam*^*KO*^ spermatids, even in cells that contained a superficially normal nucleus by phase-contrast.

**Movie S1. Sperm from *wam* mutants are completely immotile**

(A-B) Movie displaying the motility of mature sperm from *wildtype* (A) males compared to the lack of motility, which is observed in *wam*^*KO/KO*^ males (B).

## Contact for reagent and resource sharing

Further information and requests for resources and reagents should be directed to Matt Gibson.

### *Drosophila melanogaster* stocks

All stocks were maintained at 25°C on syrup food. The *wildtype* stock used was *w; His2A-mRFP;EYFP-Mito* for experiments with fluorescent imaging. *w*^*1118*^ (BDSC# 3605) was used for all other experiments.

### Generation of *wampa* transgenic lines

gRNA 1 and 2, which were used in the construction of *wam*^*KO*^, were cloned into *pCFD2-dU6.2gRNA* (Addgene, 49409) and gRNA 3, which was used for creating *wam*^*Δ1*^ and *wam*^*Δ2*^ was cloned into *pCFD3-dU6.3gRNA* (Addgene, 49410) [39]. The following primers (5’-3’) were annealed and cloned into the BbsI site:

gRNA1F: CTT CGA TAT TGG GCT TTG ATA CAC T

gRNA1R: AAA CAG TGT ATC AAA GCC CAA TAT C

gRNA2F: CTT CGG ATT TAA ATA TTC ATC AGT

gRNA2R: AAA CAC TGA TGA ATA TTT AAA TCC

gRNA3F: GTC GAA GGC GGT GTT ATC ACC ATC

gRNA3R: AAA CAT GGT GAT AAC ACC GCC TTC

To generate a donor plasmid for creating *wam*^*KO*^, we used HIFI DNA assembly master mix to assemble 4 fragments that were generated from the following primers (5’-3’):

### *w*^*1118*^ genomic DNA

(5’ homology arm F) CCT GCA GGT CGA CTC TAG AGA AGA CGG TCA TAG TGC AGG G

(5’ homology arm R) CCA AGC TTG GCG AAT AAC TTC GTA TAA TGT ATG CTA TAC GAA GTT ATA

GTG TAT CAA AGC CCA ATA TTT ATG CC

(3’ homology arm F) GTA TGT AAG TTA AAT AAC TTC GTA TAG CAT ACA TTA TAC GAA GTT ATA

CTG ATG AAT ATT TAA ATC CGT T

(3’ homology arm R) ATTCGAGCTCGGTACCCGGGTACCATCGTCGGACGTGCAT

### Ubi-mCherry DNA (Gift from Takuya Akiyama)

(mCherryF) ATA TTG GGC TTT GAT ACA CTA TAA CTT CGT ATA GCA TAC ATT ATA CGA AGT TAT

TCG CCA AGC TTG GGC TGC ATC ACG TAA TAA G

(mCherryR) GGA TTT AAA TAT TCA TCA GTA TAA CTT CGT ATA ATG TAT GCT ATA CGA AGT TAT

TTA ACT TAC ATA CAT ACT AGA ATT GAT CGG C

### pHSG98 DNA (Takara, 3298)

CCCGGGTACCGAGCTCGAATTCG

CTCTAGAGTCGACCTGCAGGCATGCAAG

The fragments were gel extracted using Zymoclean Gel DNA Recovery kit, assembled at 50°C for 3 hours, and then transformed into electrocompetent cells. The gRNA and donor templates were sequence verified, ethanol precipitated, and injected into, *y*^*2*^*cho*^*2*^*v*^*1*^; *attP40(nos-Cas9)*/+ embryos. For *wam*^*KO*^, the donor and guides 1 and 2 were mixed and injected at 250 ng/μl each (total concentration of 750 ng/μl). *wam*^*Δ1*^ and *wam*^*Δ2*^ were generated by injecting 250 ng/μl of gRNA 3. For all injections, *y2 cho2 v1; attP40(nos-Cas9)/CyO* (NIG-FLY# CAS-0001) males were crossed to *w*^*1118*^ virgins and kept at 25°C in egg laying cages on grape plates during injections and the embryos were injected at 18°C. Transformants were screened by visualization of mCherry for *wam*^*KO*^ and by homozygous male sterility for *wam*^*Δ1*^ and *wam*^*Δ2*^.

The lox-P sites that flank the m-Cherry selection cassette were removed by Cre/loxP-mediated recombination, generating the *wam*^*KO*^ line [40]. This line was recombined with *sqh-EYFP-Mito* (BDSC# 7194) and crossed to *His2Av-mRFP1* (BDSC# 23651) to generate the *w; His2Av-mRFP1 II.2/CyO*; *wam*^*KO*^, *sqh-EYFP-Mito/TM3,Sb* line that was used during fluorescent imaging analysis.

The *Wam::mScarlet-I* fusion construct and the *wam*^*SR2*^ rescue construct were created by generating a 6 kb fragment encompassing the *wampa* locus plus 2 kb on either side. The tag was placed at the C-terminus with a flexible 10x GLY linker that was incorporated into the primers. The following primers were used (5’-3’):

### CH322-169E18 (CHORI)

5’ homology arm F for tag and rescue: GCC AAG GCA AGT ATT AAA ACG T

5’ homology arm R for tag: ACC GCC TCC TCC ACC TCC GCC ACC ACC ACC TCC GCC ACC ACC ACC CAT GTT GCG CTT GGC GGC CAG GAG ACG

5’ homology arm R for rescue: TGA ATT CCG ACG GGT GCA C

3’ homology arm F for rescue: AGA TGC AGG GTA TTA TGC

3’ homology arm F for tag: TAG TTC TGG AAT ATA TGA TG

3’ homology arm R for tag and rescue: CAA CAG CAG GCG GTA TAA ATC CA

### pmScarlet-i_C1 (Addgene, 85044)

Scarlet-I F: GCG GAG GTG GAG GAG GCG GTA TGG TGA GCA AGG GCG AG

Scarlet-I R: CAT CAT ATA TTC CAG AAC TAC TTG TAC AGC TCG TCC ATG C

The fragments were gel extracted using Zymoclean Gel DNA Recovery kit, assembled using HIFI DNA assembly master mix for an hour at 50°C and transformed into electrocompetent cells. The insert was then cloned into the *w+attB* plasmid by digesting with BamHI and ApaI, for the tag construct and BamHI and NotI for the rescue construct. The ends were blunted using klenow fragment and gel purified. *w+attB* donor plasmid (Addgene, 30326) was digested with XhoI, blunted with klenow fragment, treated with shrimp alkaline phosphatase, and purified. The insert and vector were then ligated, transformed, screened, and sequence verified. Positive colonies were miniprepped and the DNA was injected into *y[1] M(vas-int.Dm)ZH-2A w[*]; PBac(y[+]-attP-3B)VK00001* (BDSC# 24861) embryos at 1:200 ng/μl. These constructs lack the 5’UTR and the first 47bp of the neighboring gene, *bb8*.

To test the functionality of the tag and rescue constructs, they were double balanced on the second and third chromosomes and crossed *wam*^*KO*^, *wam*^*Δ1*^, and *wam*^*Δ2*^. Stocks were generated that contained the either the tag or the rescue with the *wam* mutations. Homozygous males from this stock were collected and assayed for fertility. Complementation assay was performed by crossing *Mi{ET1}bb8*^*MB10362*^ (BDSC# 27841) to *wam*^*KO*^ and testing for fertility of the transheterozygotes.

### Fertility assays

For all fertility assays, 30+ single males were crossed to 3-5 *wt* (*w*^*1118*^) virgins for all indicated genotypes. Vials with progeny after 10 days were considered fertile.

### Live imaging

Sperm motility was monitored by dissecting testes of 2-3 day old *wt* and *wam*^*KO*^ males and transferring them to a clean slide in a drop of PBS. The testes were pierced to release the sperm and imaged every 2 ms for 100 frames on a Zeiss Axiovert 200M microscope using a 20x 0.8 Phase Plan-Apochromat objective.

Testis and seminal vesicle morphology were analyzed by collecting several newly eclosed males from both *wt* and *wam*^*KO*^ and holding them as virgins for two weeks. Every few days ∼5 males/genotype were analyzed. Their testes and associated seminal vesicles were dissected in PBS and transferred to a clean slide into a drop of PBS and imaged on a Leica CTR 5000 compound microscope.

Testes squashes were performed as previously described [22]. In brief, testes were dissected from 0-2 day-old *wt* (*w;His2Av,RFP;Mito,YFP*) and *wam*^*KO*^ (*w;His2Av-mRFP;wam*^*KO*^*,Mito-YFP*) homozygous males and transferred to a clean poly-L-lysine coated slide into a drop of PBS. For images showing microtubules, SiR-tubulin was added at 1:250. The testes were pierced 2-3 times to release the cells. The squashes were imaged on a Zeiss Axiovert 200M microscope using either a 20x 0.8 Phase Plan-Apochromat or 100x 1.4 Phase Plan-Apochromat objective. Brightness was occasionally adjusted for clarity and to avoid saturation.

### EM analysis

For TEM analysis, testes were dissected and fixed with a buffer containing 2.5% paraformaldehyde, 2% glutaraldehyde, 1% sucrose and 50 mM sodium cacodylate (PH7.4). After a brief rinse, the tissue was post-fixed with 1% OsO4 for 90 min and then stained with 1% uranium acetate en bloc overnight. Thereafter, the samples were dehydrated through an ethanol gradient to 100%, equilibrated with propylene oxide, and infiltrated with 50% propylene oxide/50% Epon, then 100% Epon resin (EMS, Fort Washington, PA) for 3 times over a day. After polymerizing at 60C for 48hr, the sample block was sectioned with a Leica Ultra microtome (Leica UC-6) using diamond knives. The sections were post-stained with uranyl acetate and lead citrate, and then imaged with a FEI transmission electron microscope (Tecnai Bio-TWIN 12, FEI).

### Image analysis and quantification

All analysis was performed using imageJ [41]. Brightness was adjusted as needed to avoid saturation. Area quantifications were performed on images acquired on the same scope, with the same volume of media, with the same settings using the ‘Measure’ function. The velocity analysis was performed using the ‘STICS’ (space-time image correlation spectroscopy) function as previously described [42].

Statistical analysis shown in figures 2, 3, 6, 7 were obtained using two sample t-test, *n* values are listed in the figure legends.

